# Generation of multi-transgenic pigs using PiggyBac transposons co-expressing pectinase, xylanase, cellulase, β-1.3-1.4-glucanase and phytase

**DOI:** 10.1101/2020.02.17.952515

**Authors:** Haoqiang Wang, Guoling Li, Cuili Zhong, Jianxin Mo, Yue Sun, Junsong Shi, Rong Zhou, Zicong Li, Huaqiang Yang, Zhenfang Wu, Dewu Liu, Xianwei Zhang

## Abstract

The current challenges facing the pork industry are to maximize feed efficiency and minimize fecal emissions. Unlike ruminants, pigs lack a number of digestive enzymes like pectinase, xylanase, cellulase, β-1.3-1.4-glucanase and phytase to hydrolyze the cell walls of grains to release endocellular nutrients into their digestive tracts. Herein, we synthesized multiple cellulase and pectinase genes derived from lower organisms and then codon optimized these genes to be expressed in pigs. These genes were then cloned into our previously optimized *XynB* (xylanase)- *EsAPPA* (phytase) bicistronic construct. We then successfully generated transgenic pigs that expressed four enzymes (*Pg7fn* (pectinase), *XynB* (xylanase), *EsAPPA* (phytase) and *TeEGI* (cellulase and β-glucanase)) using somatic cell cloning. Expression of these genes was parotid gland specific. Enzymatic assays using the saliva of these founders demonstrated high levels of phytase (2.0~3.4 U/mL) and xylanase (0.25~0.42 U/mL) activity, but low levels of pectinase (0.06~0.08 U/mL) activity. These multi-transgenic pigs are expected to contribute to enhance feed utilization and reduce environmental impact.

## Introduction

In the pig industry, ineffective digestion causes excess nutrients to be released to the environment. This results in soil salinity and potential pollution to water and air[1]. Domestic pigs mainly feed on common cereal grains, oil seed meals and their by-products. These contain various anti-nutritional factors such as non-starch polysaccharides and phytic acid[2,3]. These anti-nutritional factors have an obvious effect on the digestion and absorption of nutrients. This is because it hinders the contact of endogenous digestive enzymes with chyme and hence slows down the nutritional diffusion rate into the intestines[1]. As a consequence, undigested nutrients containing large amounts of inorganic nitrogen and phosphorus are excreted by the pigs to stimulate growth of algae and other aquatic plants and hence enhance microbial proliferation that ultimately contributes to air pollution.

Several dietary manipulation strategies have been employed to reduce fecal output and nutrient excretion in swine. The most widely practiced strategy is to introduce phytate- or non-starch polysaccharides- degrading enzymes in the formula feed. These can effectively decrease nitrogen and/or phosphorus emissions and hence reduce environmental impact. However, various factors affect the catalytic activity of these microbial enzymes, such as feed processing and storage, feed components, pH, minerals and temperature. Recently, genetically engineered pigs that express specific or multiple digestive enzyme genes have provided an alternative strategy to replace dietary enzyme supplementation in feed. Recently study demonstrated that transgenic pigs that produce salivary phytase had less than 75% of fecal phosphorus and required almost no inorganic phosphate supplementation for normal growth compared to non-transgenic pigs[4]. In our previous study, we established transgenic pigs that simultaneously expressed three microbial enzymes, β-glucanase, xylanase, and phytase in their salivary glands. This significantly enhanced growth and reduced fecal nitrogen and phosphorus levels in pigs[5].

In this study, we isolated and characterized several novel digestive enzyme genes, and then generated transgenic pigs that expressed these multiple enzymes, like pectinase, xylanase, cellulase, β-1.3-1.4-glucanase and phytase. These genes were expressed using a salivary gland promoter. The transgenic pigs had no adverse reactions and had better feed digestion compared to non-transgenic pigs.

## Results

### Characterization of the three pectinase genes expressed in PK-15 cells

Based on a previous study, we initially selected three pectinase genes *Pg7fn*, *PgaA* and *PGI* for our studies. Enzyme activity assays demonstrated that *Pg7fn* had the highest pectinase activity towards 1% polygalacturonic acid and 55%~70% for esterified pectin as the substrates, respectively. *PGI* had the second highest pectinase activity towards 1% polygalacturonic acid. However, the activity of *Pg7fn*, *PgaA* and *PGI* was less than 0.1 U/mL for > 85% esterified pectin **(Fig 1a, b and c)**. We selected *Pg7fn* and *PGI* to determine their optimal pH in 1% polygalacturonic acid.

**Fig 1.**
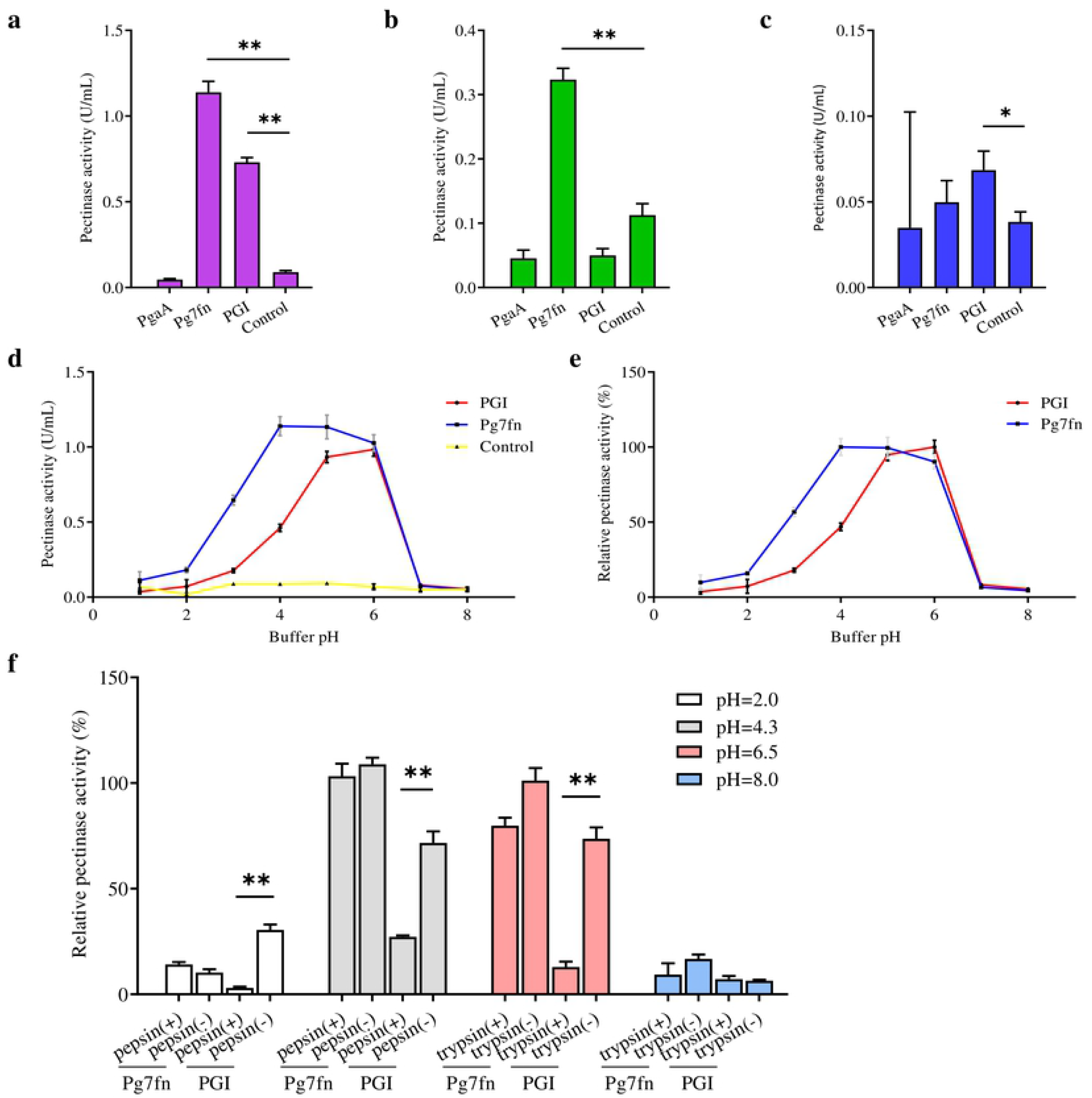
Characterization of the three pectinase genes expressed in PK-15 cells. Pectinase activities of *PgaA, Pg7fn* and *PGI* were evaluated using **(a)** 1% poly-galacturonic acid, **(b)** 55%~70% esterified pectin and **(c)** > 85% esterified pectin as substrates at pH 4.5, respectively. **(d)** Pectinase activity and **(e)** relative pectinase activity of *Pg7fn* and *PGI* at different pH levels (1.0~8.0). **(f)** *Pg7fn* and *PGI* were incubated with different pepsin and trypsin pH solutions at 39.5°C for two hours. Control represents pcDNA 3.1(+) vector. Data is shown as mean ± SEM, n = 3 (one-way ANOVA). * *P* < 0.05, ** *P* < 0.01.

Enzyme activity of *Pg7fn* increased with pH between 1.0~4.0 and reached highest pectinase activity at pH 4.0, at approximately 1.15 U/mL. The high enzyme activity was stable at pH 4.0~6.0, and then decreased significantly after pH 6.0. *PGI* showed the same trend with *Pg7fn*, but reached its highest enzyme activity at pH 6.0 **(Fig 1d)**. The relative pectinase activity of *Pg7fn* and *PGI* remained at least 56.8% and 46.8% during the stationary phase, respectively **(Fig 1e)**. To simulate the pig’s digestive tract, we treated *Pg7fn* and *PGI* at 39.5°C for two hours with different pepsin and trypsin pH solutions. The results indicated that pectinase activity of *PGI* was significantly decreased after pepsin or pH 6.5 trypsin treatment **(Fig 1f)**. However, *Pg7fn* was not affected by treatment with pepsin and trypsin. Hence, *Pg7fn* was selected as the candidate gene.

### Characterization of the six cellulase genes expressed in PK-15 cells

We selected six endo-β-1,4-endoglucanase genes *cel5B*, *egII*, *AG-egaseI*, *TeEGI*, *cel9* and *Bh-egaseI* to measure cellulase and β-glucanase activity at various pH conditions. *egII* and *TeEGI* cellulase activity were significantly higher (0.27 U/mL and 0.28 U/mL, respectively **(Fig 2a)**) compared to the other genes for 1% sodium carboxymethyl cellulose. Furthermore, β-glucanase activity of *egII* and *TeEGI* were approximately 0.76 U/mL and 0.86 U/mL for 0.8% β-D-glucan as substrate, respectively. The other genes had activities of less than 0.09 U/mL **(Fig 2b)**. To further clarify the enzymatic characteristics of *egII* and *TeEGI*, we optimized the pH levels of the reaction buffer. We found that *TeEGI* had the highest cellulase activities at pH 4.5 and had high residual activity after treatment with pH 3.5~7.0 **(Fig 2c and d)**. *egII* had similar trends, however the optimal pH was 5.0. The β-glucanase activity of *TeEGI* was greater than 0.88 U/mL at pH 3.0~7.0 and reached the maximum of 1.11 U/mL at pH 5.5 **(Fig 2e and f)**. Compared to *TeEGI*, the highest β-glucanase activity of *egII* was 0.77 U/mL and had residual activity of greater than 50% between pH 2.0~7.0. We then investigated whether *egII* and *TeEGI* would have high enzyme activity in different pepsin and trypsin pH buffers. The results indicated that *TeEGI* was resistant to pepsin and trypsin digestion, but *egII* β-glucanase and cellulase were significantly inhibited at pH 2.0 pepsin buffer **(Fig 2g and h)**. Hence, we selected *TeEGI* as the candidate cellulase and β-glucanase gene.

**Fig 2.**
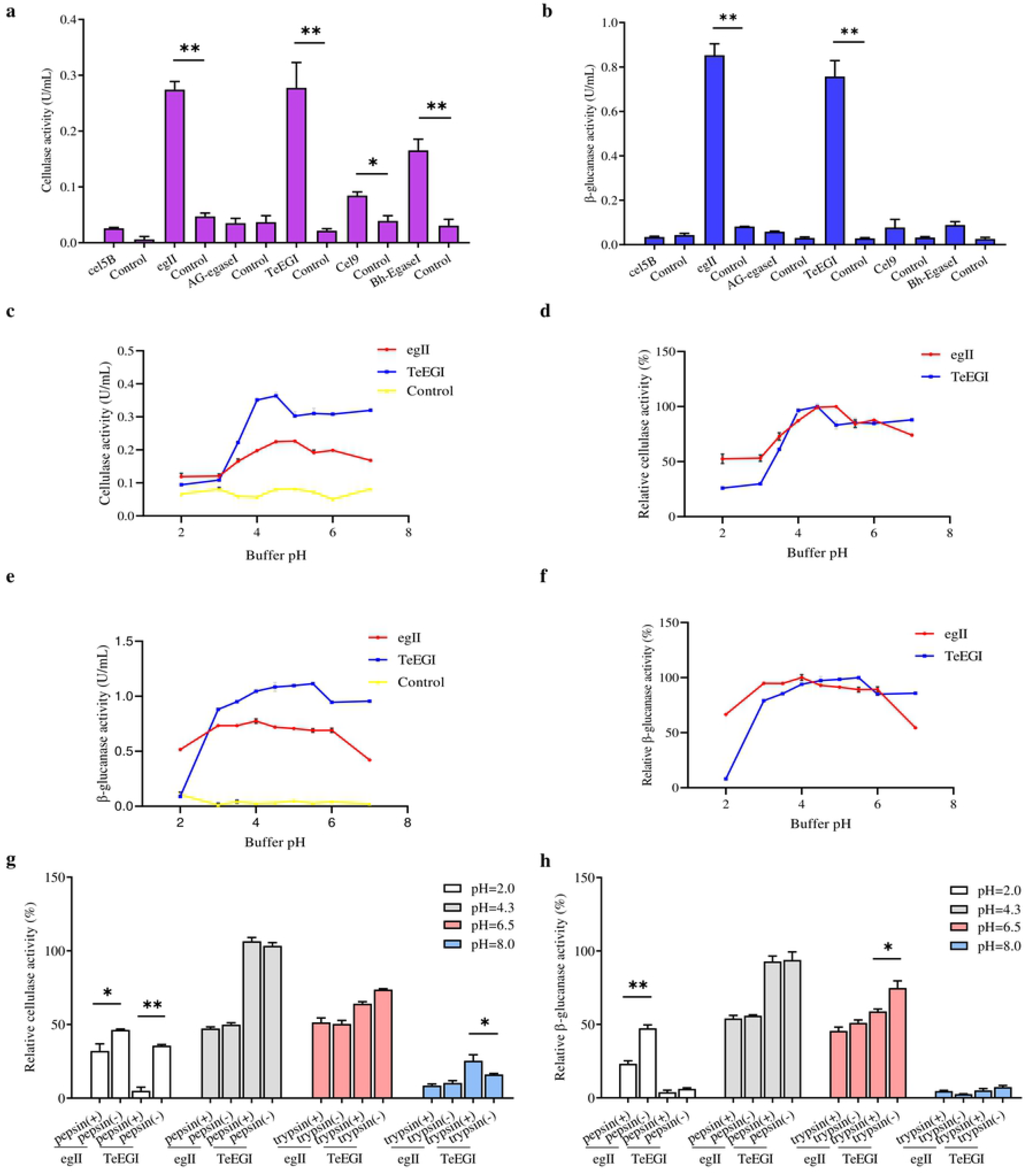
Characterization of six cellulase genes expressed in PK-15 cells. **(a)** cellulase or **(b)** β-glucanase activities of *cel5B*, *egII*, *AG-egaseI*, *TeEGI*, *cel9* and *Bh-egaseI* were evaluated at suitable pH conditions. **(c)** Cellulase activity and **(d)** relative activity of *egII* and *TeEGI* at different pH levels (2.0~7.0). **(e)** β-glucanase activity and **(f)** relative activity of *egII* and *TeEGI* at different pH levels (2.0~7.0). **(g)** Cellulase and **(h)** β-glucanase activity of *egII and TeEGI* were measured following incubation with different pepsin and trypsin pH solutions. Control represents pcDNA3.1(+) vector. Data is shown as mean ± SEM, n = 3 (t-test). * *P* < 0.05 or ** *P* < 0.01.

### Enzyme activity between polycistronic and monomeric constructs

To assess the polycistronic positions of the four genes (*Pg7fn*, *TeEGI*, *EsAPPA* and *xynB*), we initially included the 2A linker at the end of each corresponding gene. Previous studies had demonstrated that XynB protein with P2A residue at the C-terminus still had high xylanase activity in porcine saliva^5^. Our results demonstrated that the enzymatic activities of EsAPPA and Pg7fn with 2A residue also kept high relative activity (> 77% and > 92%, respectively) **(Fig 3a and b)**. However, cellulase and β-glucanase activity of *TeEGI* with T2A was significantly reduced to 64.8% and 55.1%, respectively **(Fig 3c)**. We fused *Pg7fn*, *XynB*, *EsAPPA* and *TeEGI* genes head to tail with E2A, P2A and T2A linkers, and named the final construct *PXAT* **(Fig 3d)**. *PXAT* was then ligated into pcDNA3.1(+) to evaluated enzyme activity. The results showed that using *PXAT,* the pectinase, xylanase, phytase, cellulase and β-glucanase enzyme activities were significantly reduced to 31.0%, 23.5%, 30.2%, 24.5% and 24.4%, respectively, compared to constructs expressing a single gene **(Fig 3e)**. mRNA levels further confirmed that the four genes that were co-expressed were lower compared to mRNA levels expressed by the single gene constructs **(Fig 3f)**.

**Fig 3.**
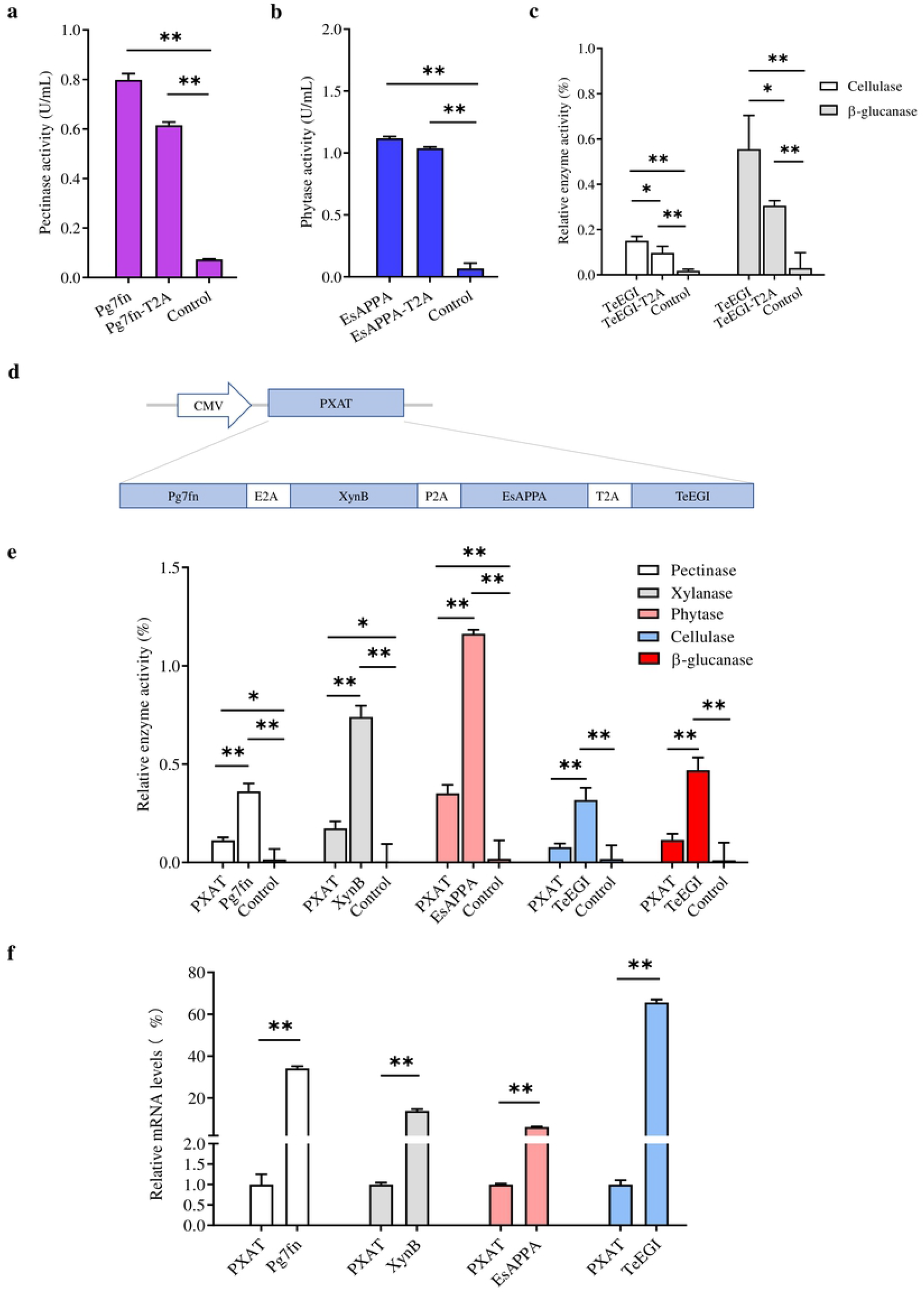
Enzyme activity between the polycistronic and single gene vector construct. The effect of 2A linker peptide on **(a)** pectinase, **(b)** phytase, **(c)** cellulase and β-glucanase activity. **(d)** Schematic of the *PXAT* vector. **(e)** Enzyme activity between PXAT and its corresponding protein expressed by the single gene constructs. **(f)** relative mRNA expression levels between genes expressed with PXAT and single gene constructs. Control represents pcDNA3.1(+) vector. Data is shown as mean ± SEM, n = 3 (one-way ANOVA). * *P* < 0.05 or ** *P* <0.01.

### Generation and identification of transgenic pigs

*PXAT* was also inserted into the tissue-specific vector pPB-mPSP-loxp-neoEGFP-loxp to form the final transgene construct (mPSP-PXAT) **(Fig 4a)**. The mPSP-PXAT contained the mouse parotid secretory protein (mPSP) promoter, loxp flanking the neo-EGFP marker genes and the left and right ends of the PiggyBac elements. For transgene cell line selection, PFFs were co-electroporated and G418 was used for selection. The EGFP marker gene was deleted in clonal cells using Cre enzyme prior to somatic cell nuclear transfer **(Fig 4b)**. A total of two cell lines were pooled and used as nuclear donors. We transferred a total of 2,096 reconstructed embryos into 10 recipient gilts. Four recipients became pregnant and delivered 9 Duroc piglets, of which 7 were alive and 2 were dead **(S1 Table)**. PCR sequencing demonstrated that 5 founders were positive for the transgene **(Fig 4c)**, but only 3 of which were alive **(Fig 4d)**. Southern blot and quantitative PCR demonstrated that three piglets carried two copies of the transgene **(Fig 4e and f)**. A positive boar was euthanized and tissue samples collected to determine expression levels of transgenic mRNA at 10 months of age. The results showed that the four genes, i.e., *Pg7fn*, *XynB*, *EsAPPA* and *TeEGI* were highly expressed in the parotid gland, had low expression in the sublingual and submandibular gland, and not expressed in other tissues **(S1 Fig)**. Enzymatic activity assays showed that the saliva from three founders were positive for pectinase, xylanase, and phytase (0.06~0.08 U/mL, 0.24~0.42 U/mL, 1.9~3.4 U/mL, respectively) **(Fig 4h~l)**. Interestingly, although we were unable to detect cellulase and β-glucanase activity, the western blotting analysis indicated that the four genes (*PXAT*) were expressed **(Fig 4g)**. The F1 pigs were obtained from 920307 transgenic pig by mating with 2 wild-type gilts, the results revealed that F1 pigs had pectinase, xylanase and phytase activities, but no cellulase and β-glucanase activities **(S2 Fig)**, which were consistent with the founders. The growth rate of F1 transgenic pigs and wild-type littermates were measured, which shown that PXAT pigs had a tendency to improve growth performance. It took an average of 84 days for transgenic pigs to grow from 30 to 100kg, whereas wild-type pigs required about 96 days **(S3 Fig)**. We also measured serum biochemical markers in both F1 transgenic and wild-type pigs **(S2 Table)**. The results showed that the phosphorus content of transgenic pigs (3.32 mM) was higher compared to wild-type pigs (2.79 mM).

**Fig 4.**
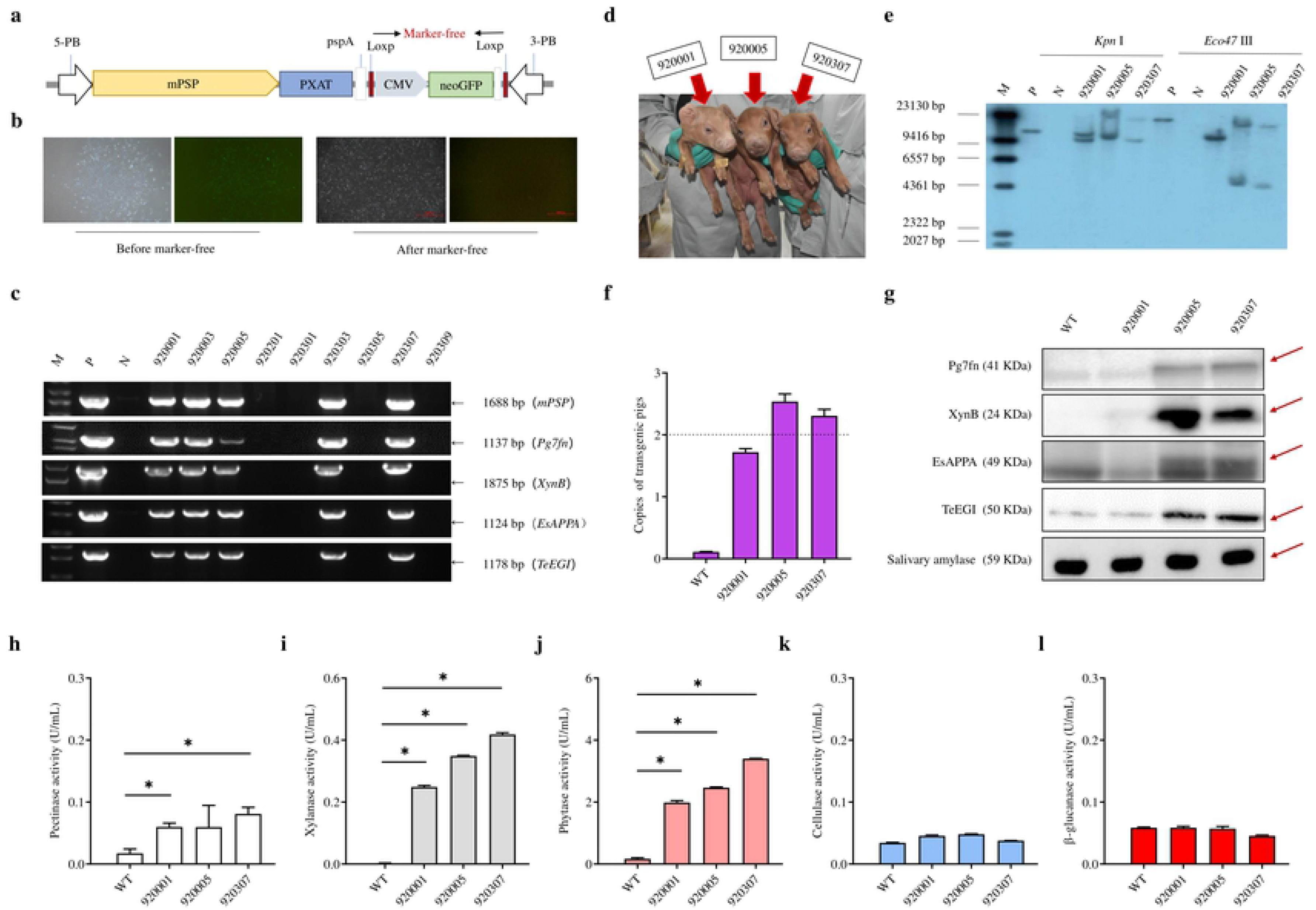
Generation and identification of the transgenic pigs. **(a)** Schematic of the transgenic plasmid mPSP-PXAT. The mPSP-PXAT consisted of the mouse parotid secretory protein (mPSP) promoter, loxp system with the neo-EGFP marker protein, and a PiggyBac transposon. **(b)** EGFP was deleted using Cre recombinase prior to somatic cell nuclear transfer. **(c)** Genomic identification of transgenic piglets using PCR and gel electrophoresis. **(d)** Transgenic piglets at 2 weeks old (Original image in **S2 Raw image**). **(e)** Southern blot analysis of transgene integration in transgenic piglets. Genomic DNA was digested using *Kpn* I and *Eco47* III endonucleases. **(f)** Copy number determination in transgenic piglets by absolute quantification. **(g)** Western blotting analysis of Pg7fn, XynB, EsAPPA and TeEGI protein expression. Salivary amylase was used as a protein reference. **(h)** Salivary pectinase, **(i)** xylanase, **(j)** phytase, **(k)** cellulase and **(l)** β-glucanase expression at 4 months. The irrelevant lanes of gels and blots were removed (Images in **S1 Raw images**). M is the DNA marker, P indicates mPSP-PXAT plasmid; N and WT represent wild-type pigs. Data is shown as mean ± SEM (one-way ANOVA). * *P* < 0.05.

## Discussion

Environmentally-friendly transgenic pigs could efficiently improve the absorption of anti-nutritional factors, enhance their growth, and reduced the emission of nitrogen and phosphorus to the environment[5]. Previous studies have demonstrated that salivary phytase and xylanase produced from transgenic pigs could effectively reduce phosphorus and nitrogen emissions[4,6]. However, no studies to date have investigated cellulase or pectinase transgenic pigs. In this study, we initially selected three pectinase genes (*PgaA, Pg7fn and PGI*) and six cellulase genes (*cel5B, egII, AG-egaseI, TeEGI, cel9* and *Bh-egaseI*) based on previous studies. Our results demonstrated that *Pg7fn* and *TeEGI* had high enzyme activity at different pH levels, and maintained their stability in different pepsin and trypsin pH buffers. However, several genes had no detectable enzyme activity. These genes were derived from microorganisms and insects, and the PK-15 cell line that was used to express these genes were unable to properly recapitulate the post-translation modifications needed for enzyme activity. In addition, the polycistronic order of the four genes (*Pg7fn*, *XynB*, *EsAPPA* and *TeEGI*) were constructed using the 2A linker at the end of each corresponding gene. In PK-15 cells, our result demonstrated that the target protein with 2A residue at the C-terminus significant reduced enzyme activity, such as Pg7fn, EsAPPA and TeEGI, in which, the activity of cellulase and β-glucanase (TeEGI) decline is most pronounced. It seemed that particular protein require special folding compared to the others. The 2A linker is derived from viruses, such as foot-and-mouth- disease virus (F2A), equine rhinitis A virus (E2A), thosea asigna virus (T2A), and porcine teschovirus-1 (P2A). When mRNA is translated, ribosomes jump from Gly to Pro in the 2A sequence. This results in the absence of a peptide bond between Gly and Pro. As a consequence, the upstream protein that is generated has a 17~19 amino acid peptides that contains Gly at the C-terminus, while the downstream protein that is generated has a Pro residue at the N-terminus, which may affect the spatial folding of the protein. As mentioned previously, the incomplete cleavage of the 2A linker could reduce protein expression[7]. There is a parotid gland expression signal peptide in front of each gene, which seems to rule out the reason that the C-terminal protein stays on the endoplasmic reticulum due to the inability of 2A linker to completely cleave[8]. In addition, it is possible that some proteins are unable to be completely synthesized due to incomplete translation. This may explain why some of the PXAT enzyme activities were significantly reduced compared to proteins that were synthesized using the single-gene vector. Finally, the larger size of the PXAT construct may contribute to lower transfection efficiency compared to constructs having only a single gene[9].

We successfully generated three transgenic pigs expressing multiple digestive enzyme genes using the PiggyBac transposon system. Although the transgenic pigs could efficiently express pectinase, xylanase and phytase, we were unable to detect the enzyme activity of cellulase and β-glucanase. Western blot analysis indicated that the TeEGI protein was expressed. Previous study suggested that different post-translational modification manners have an effect on protein function[10]. Thus, we suspect that TeEGI lacks cellulase activity, possibly due to post-translational modification that alters the folding or function of the protein. Additional, although cellulase was secreted in PK-15 cells, interaction between various cells in an individual could also affect protein function. The polyA tail plays a crucial role in transcription, translation and stabilization of mRNAs[11]. In our previous work, we used the *bGH-pA* (bovine growth hormone polyadenylation signal) as a termination sequence[5]. But in this study, we firstly utilized an unconventional polyA (3’ UTR of parotid secretory protein as a termination sequence *pspA*) in order to evaluate its effect **(Fig 4a)**. We infered that *pspA* may affect the activity of cellulase and β-glucanase. Due to the unavailability of porcine parotid gland cell lines, we used the PK-15 cell line to express the four enzyme genes driven by the CMV promoter. However, in animal models, multiple digestive enzyme genes are driven by the parotid secretory protein promoter. Hence, the low enzyme activity that was observed may be due to an incompatible promoter.

In summary, we successfully produced transgenic pigs using somatic cell transfer. These transgenic pigs expressed, under the control of parotid gland specific promoter, four enzyme genes (*Pg7fn* (pectinase), *XynB* (xylanase), *EsAPPA* (phytase) and *TeEGI* (cellulase and β-glucanase)). These transgenic animals are expected to offer a valuable experience for the global environmental concerns and the inefficient absorption of feed in livestock.

## Materials and Methods

### Plasmid construction

Three pectinase genes, *PgaA* (*Aspergillus niger* JL-15)[12], *Pg7fn* (*Thielavia arenaria* XZ7)[13] and *PGI* (*chaetomium sp*)[14]; one xylanase *gene XynB* (*Aspergillus niger*)[5,15], one phytase gene *EsAPPA* (*Escherichia coli*)[5] and six cellulase and β-glucanase genes (respectively), *cel5B* (*Gloeophyllum trabeum*)[16], *egII* (*Pichia pastoris*)[17], *AG-egaseI* (*Apriona germari*)[18], *TeEGI* (*Teleogryllus emma*)[19], *cel9* (*Clostridium phytofermentans*)[20] and *Bh-egaseI* (*Batocera horsfieldi*)[21] were optimized and synthesized based on pig codon preferences using Genscript (Nanjing, China). They were then cloned into pcDNA3.1(+). *Pg7fn*, *XynB*, *EsAPPA* and *TeEGI* genes were then head-to-tail ligated using E2A, P2A and T2A linkers. The ligated construct was named *PXAT*. *PXAT* was then inserted into pcDNA3.1(+) and enzyme activity was evaluated. *PXAT* was also inserted into the tissue-specific vector pPB-mPSP-loxp-neoEGFP-loxp[5] to generate the final transgene construct (mPSP-PXAT). The primer sets used for cloning are listed in **S3 Table**.

### Cell culture and transfection

The PK-15 cell line (ATCC CCL-33) and porcine fetal fibroblasts (PFFs) were cultured in DMEM (Thermo Fisher Scientific, Suwanee, GA,USA) supplemented with 10% fetal bovine serum (Thermo Fisher Scientific, Suwanee, GA,USA). To evaluate enzyme activity, PK-15 were grown to 70% confluence, and then transfected using lipofectamine LTX reagent (Thermo Fisher Scientific, Suwanee, GA,USA). 60 hrs post-transfection, the culture supernatant was collected for enzyme assays. For transgene cell line selection, PFFs were co-electroporated with a circular transposase plasmid pCMV-hyPBase and a circular mPSP-PXAT plasmid using the program A-033 on the Nucleofector 2b Device (Amaxa Biosystems/Lonza, Cologne, Germany). After cell attachment, 400 µg/ml G418 (Gibco) was added to the culture media for cell selection. Clonal cells expressing green fluorescence were selected and identified by PCR and sequencing.

### Generation of transgenic pigs

The EGFP marker gene and neomycin resistant gene (neoR) were removed from transgenic cells using Cre enzyme (Excellgen, Rockville, MD USA) and then mixed multiple positive clones as nuclear donors for somatic cell nuclear transfer. Somatic cell nuclear transfer was described as previously studied[5]. The reconstructed embryos were transferred into recipient gilts, and piglets were naturally born after gestation. Afterwards, genomic DNA was extracted and sequenced using PCR (**S4 Table**). Additionally, mRNA was extracted from porcine tissue samples and reversed transcribed to cDNA to be used as the template for qPCR. Relative quantitative Real-time PCR was used to identify mRNA expression levels in transgenic pigs and absolute quantitative Real-time PCR was used to detect copy number in transgenic pigs (primers used are listed in **S5 Table**).

### Southern and western blot analysis

Genomic DNA was digested with restriction enzymes *Kpn* I or *Eoc47* III, and then run on an 0.8% agarose gel. The digested fragments were then transferred to a nylon membrane. The membrane was hybridized using digoxigenin-labeled DNA probes (**S4 Table**) for *mPSP* based on the DIG-High Prime DNA Labeling and Detection Starter Kit II protocol (Roche, Mannhein, Germany). For western blotting, saliva was collected and then ultra-filtrated using a centrifugal filter (Millipore, Massachusetts, USA). Total protein from saliva was then electrophoresed on an SDS polyacrylamide gel, and subsequently transferred to a polyvinylidene fluoride membrane (Millipore, Massachusetts, USA). The membranes were incubated overnight at 4°C with primary rabbit polyclonal antibodies (**S6 Table**) against Pg7fn, XynB, EsAPPA or TeEGI (purchased from Genscript, Nanjing, China). The salivary amylase antibody (ab34797, Abcam) was used to confirm equal protein loading and the dilution ratio was 1: 1000. Membranes were then washed and incubated with a secondary IgG antibody. Bands were visualized using the UVP software.

### Enzyme analysis assay

Cell culture supernatants and porcine saliva were centrifuged and used for enzyme analysis assays. Pectinase, xylanase, β-glucanase and cellulase activity were assayed using 1% (w/v) polygalacturonic acid (and 55%~70% esterified pectin, > 85% esterified pectin), 1% (w/v) xylan, 0.8% (w/v) β-D-glucan, and 1% (w/v) sodium carboxymethyl cellulose as the substrates, respectively. Reducing sugar content was measured using the 3,5-dinitrosalicylic acid (DNS) method[5,14,16]. One unit of enzyme activity was defined as the rate at which 1 μmol of reducing sugar was released per minute. Phytase activity in saliva was measured as previously described[5].

The optimal pH of these proteins were determined at 39.5°C for 30 min in buffers of pH 1.0~8.0. The buffers used were 0.2 M potassium chloride (KCl)- hydrochloric acid (HCl) for pH 1.0, 0.2 M glycine-HCl for pH 2.0~3.0, and 0.2 M citric acid-disodium hydrogen phosphate (Na_2_HPO_4_) for pH 4.0~8.0. All protein tolerance tests were measured after buffer treatment for 2 h under optimal conditions (optimal pH, 39.5°C and 30 min).

### Feeding management

Transgenic pigs and wild-type littermates were fed on the same diet (**S7 Table**). They were raised in the same pens fitted with MK3 FIRE feeders (FIRE, Osborne Industries Inc., Osborne, KS). Individual daily feed intake and body weights were recorded when the pigs accessed the FIRE feeders. All pigs had free access to feed and drinking water throughout the growth phase. Blood was sterile collected at 90 days of age. Serum biochemical parameters of growing-finishing pigs were determined using a Hitachi 7020 full-automatic biochemical analyzer (Japan).

### Statistical analysis

Data was analyzed using the IBM SPSS Statistics 20 (IBM SPSS, Chicago, IL, USA) or SAS9.4 (SAS Inst. Inc., Cary, NC, USA). For enzyme analysis and relative gene expression, one-way ANOVA was used. For serum biochemical data, unpaired t-test (two-tailed) was used. For growth performance, a total of 3 F1 transgenic pigs (1 boar, 2 gilts) and 6 wild-type littermates (3 boars, 3 gilts) were test. When it comes to statistics, multivariate analysis of variance (MANOVA) was performed using the GLM procedure, with sex and initial weight used as the covariate. Data was expressed as mean ± SEM. *P* < 0.05 considered statistically significant.

## Acknowledgments

We gratefully acknowledge Ranbiao Mai, Wanxian Yu, and Lvhua Luo (Wens Foodstuff Group Co., Ltd) for somatic cell nuclear transfer and Wenxu Feng for embryo transfer. We thank Xiaofang Ruan and Guangyan Huang (South China Agricultural University) for saliva and tissue sample collection.

## Supporting information

**S1 Raw images. The full scan of all gels and blots**.

**S2 Raw image. The raw image of transgenic piglets at 2 weeks old**.

**S1 Fig. Relative mRNA levels of *PXAT* in various tissues from 10-month old transgenic pigs**.

**S2 Fig. The enzyme activities of the F1 transgenic pigs**.

**S3 Fig. Growth rate of F1 transgenic pigs and wild-type littermates during the growing period (30 kg to 100 kg)**.

**S1 Table. Embryo transfer data for cloned pigs**.

**S2 Table. Serum biochemical information for F1 transgenic pigs.**

**S3 Table. Primers used in vector construction**.

**S4 Table. Primers used in PCR and probes in southern blotting**.

**S5 Table. Primers used in quantitative real-time PCR and absolute quantitative real-time PCR**.

**S6 Table. Customized primary antibody information in western blotting**.

**S7 Table. Pig growth and fattening stage feed formulas**.

## References

1. Shirali M, Doeschl-Wilson A, Knap PW, Duthie C, Kanis E, van Arendonk JA, et al. Nitrogen excretion at different stages of growth and its association with production traits in growing pigs. J Anim Sci. 2012; 90(6): 1756–65. PMID: 22178856.

2. Gilani GS, Cockell KA, Sepehr E. Effects of antinutritional factors on protein digestibility and amino acid availability in foods. J AOAC Int. 2005; 88(3): 967–87. PMID: 16001874.

3. Bohn L, Meyer AS, Rasmussen SK. Phytate: impact on environment and human nutrition. A challenge for molecular breeding. J Zhejiang Univ Sci B. 2008; 9(3): 165–91. PMID: 18357620.

4. Golovan SP, Meidinger RG, Ajakaiye A, Cottrill M, Wiederkehr MZ, Barney DJ, et al. Pigs expressing salivary phytase produce low-phosphorus manure. Nat Biotechnol. 2001; 19(8): 741–5. PMID: 11479566.

5. Zhang X, Li Z, Yang H, Liu D, Cai G, Li G, et al. Novel transgenic pigs with enhanced growth and reduced environmental impact. Elife. 2018 May 22; 7. pii: e34286. PMID: 29784082.

6. Zhang M, Cai G, Zheng E, Zhang G, Li Y, Li Z, et al. Transgenic pigs expressing beta-xylanase in the parotid gland improve nutrient utilization. Transgenic Res. 2019; 28(2): 189–98. PMID: 30637610.

7. Velychko S, Kang K, Kim SM, Kwak TH, Kim KP, Park C, et al. Fusion of Reprogramming Factors Alters the Trajectory of Somatic Lineage Conversion. Cell Rep. 2019; 27(1): 30–9. PMID: 30943410.

8. de Felipe P, Luke GA, Brown JD, Ryan MD. Inhibition of 2A-mediated ‘cleavage’ of certain artificial polyproteins bearing N-terminal signal sequences. Biotechnol J. 2010; 5(2): 213–23. PMID: 19946875.

9. Kreiss P, Cameron B, Rangara R, Mailhe P, Aguerre-Charriol O, Airiau M, et al. Plasmid DNA size does not affect the physicochemical properties of lipoplexes but modulates gene transfer efficiency. Nucleic Acids Res. 1999; 27(19): 3792–8. PMID: 10481017.

10. Knorre DG, Kudryashova NV, Godovikova TS. Chemical and functional aspects of posttranslational modification of proteins. Acta Naturae. 2009; 1(3): 29–51. PMID: 22649613.

11. Edmonds M. A history of poly A sequences: from formation to factors to function. Prog Nucleic Acid Res Mol Biol. 2002; 71: 285–389. PMID: 12102557.

12. Liu MQ, Dai XJ, Bai LF, Xu X. Cloning, expression of *Aspergillus niger JL-15* endo-polygalacturonase A gene in *Pichia pastoris* and oligo- galacturonates production. Protein Expr Purif. 2014; 94: 53–9. PMID: 24231374.

13. Tu T, Meng K, Huang H, Luo H, Bai Y, Ma R, et al. Molecular characterization of a thermophilic endopolygalac-turonase from *Thielavia arenaria XZ7* with high catalytic efficiency and application potential in the food and feed industries. J Agric Food Chem. 2014; 62(52): 12686–94. PMID: 25494480.

14. Tu T, Meng K, Bai Y, Shi P, Luo H, Wang Y, et al. High-yield production of a low-temperature-active polygalacturonase for papaya juice clarification. Food Chem. 2013; 141(3): 2974–81. PMID: 23871048.

15. Deng P, Li DF, Cao YH, Lu WQ, Wang CL. Cloning of a gene encoding an acidophilic endo-beta-1,4-xylanase obtained from *Aspergillus niger* CGMCC1067 and constitutive expression in *Pichia pastoris*. Enzyme Microb Technol. 2006; 39(5): 1096–102. https://doi.org/10.1016/j.enzmictec.2006.02.014.

16. Kim HM, Lee YG, Patel DH, Lee KH, Lee DS, Bae HJ. Characteristics of bifunctional acidic endoglucanase (*Cel5B*) from *Gloeophyllum trabeum*. J Ind Microbiol Biotechnol. 2012; 39(7): 1081–9. PMID: 22395898.

17. Akbarzadeh A, Ranaei Siadat SO, Motallebi M, Zamani MR, Barshan Tashnizi M, Moshtaghi S. Characterization and high level expression of acidic endoglucanase in *Pichia pastoris*. Appl Biochem Biotechnol. 2014; 172(4): 2253–65. PMID: 24347161.

18. Lee SJ, Kim SR, Yoon HJ, Kim I, Lee KS, Je YH, et al. cDNA cloning, expression, and enzymatic activity of a cellulase from the mulberry longicorn beetle, *Apriona germari*. Comp Biochem Physiol B Biochem Mol Biol. 2004; 139(1): 107–16. PMID: 15364293.

19. Kim N, Choo YM, Lee KS, Hong SJ, Seol KY, Je YH, et al. Molecular cloning and characterization of a glycosyl hydrolase family 9 cellulase distributed throughout the digestive tract of the cricket *Teleogryllus emma*. Comp Biochem Physiol B Biochem Mol Biol. 2008; 150(4): 368–76. PMID: 18514003.

20. Zhang XZ, Sathitsuksanoh N, Zhang YH. Glycoside hydrolase family 9 processive endoglucanase from *Clostridium phytofermentans*: Heterologous expression, characterization, and synergy with family 48 cellobiohydrolase. Bioresour Technol. 2010; 101(14): 5534–8. PMID: 20206499.

21. Mei HZ, Xia DG, Zhao QL, Zhang GZ, Qiu ZY, Qian P, et al. Molecular cloning, expression, purification and characterization of a novel cellulase gene (*Bh-EGaseI*) in the beetle *Batocera horsfieldi*. Gene. 2016; 576: 45–51. PMID: 26410410.

